# Maftools: Efficient analysis, visualization and summarization of MAF files from large-scale cohort based cancer studies

**DOI:** 10.1101/052662

**Authors:** Anand Mayakonda, H Phillip Koeffler

## Abstract

Mutation Annotation Format (MAF) has become a standard file format for storing somatic/germline variants derived from sequencing of large cohort of cancer samples. MAF files contain a list of all variants detected in a sample along with various annotations associated with the putative variant. MAF file forms the basis for many downstream analyses and provides complete landscape of the cohort. Here we introduce maftools–an R package that provides rich source of functions for performing various analyses, visualizations and summarization of MAF files. Maftools uses data.table library for faster processing/summarization and ggplot2 for generating rich and publication quality visualizations. Maftools also takes advantages of S4 class system for better data representation, with easy to use and flexible functions.

**Availability and Implementation:** maftools is implemented as an R package available at https://github.com/PoisonAlien/maftools

## Introduction

With advances in cancer genomics and reduction in cost per base of sequencing technologies, sequencing large cohort of cancer patients has become an efficient way of determining genetic abnormalities associated with the disease [1–4]. Such cohort-based studies often results in large amount of data in the form of somatic/germline variants containing single nucleotide variants (SNP) and small insertion/deletions (INDELS). This data is generally stored in the form of Mutation Annotation Format and provides a complete genomic landscape of the cohort [5]. The Cancer Genome Atlas (TCGA) project has sequenced over 30 different types of cancer and resulting somatic variants are stored as MAF files, with several independent studies following the same [6].

MAF files provides baseline data for many downstream analyses such as driver gene detection, detecting mutually exclusive set of events, mutational signatures and tumor heterogeneity estimation [7–10]. Visualization also plays key role in genomic studies, with researchers often struggling to generate publication quality images, such as oncoplots (also known as waterfall plots), lollipop plots and oncoprints to name a few. As MAF files are getting standardized, current bioinformatic community lacks software to process them. Here, we developed maftools to process, summarize and analyze MAF files, resulting from large cohort based studies. Maftools provides various plotting functions to visualize data stored in MAF files to help researchers generate publication quality images. Functions are also implemented to perform some of the common analyses in cancer studies, including disease associated driver gene detection, mutual exclusivity analysis, and tumor heterogeneity estimation. Along with analysis of MAF files, maftools also provides functions to integrate and visualize of copy number data. Usage of maftools is straightforward with self-explanatory functions and is implemented as an open source R package.

Functions implemented in maftools can be broken down into four main categories as shown in (Figure 1.

**Figure 1.**
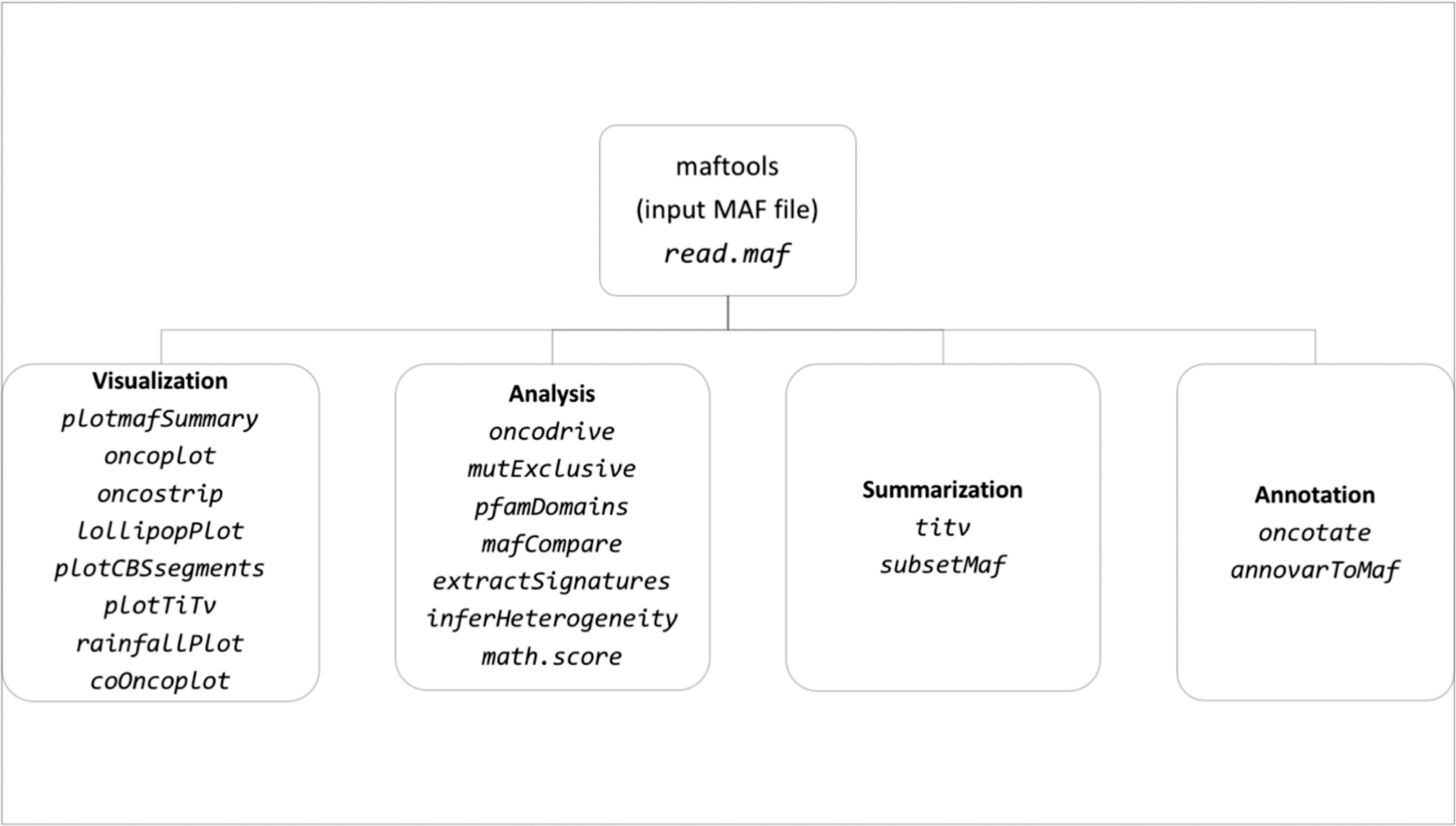
List of available functions in maftools, classified into four main categories. Function names are in *italics*.

Here we briefly demonstrate the application of maftools on TCGA acute myeloid leukemia cohort (LAML) [11]. All of the raw data and code used to generate following results are included in the installation package and demonstrated as a vignette.

## Visualization

- MAF summary and Oncoplots: Oncoplots are frequently used in cohort based cancer studies to display complete mutational landscape. Oncoplots consist of main matrix with each row representing a gene and each column representing a sample. Matrix is organized and sorted to display frequently mutated genes at the top. Occasionally these plots include top and side bar plots to show numbers associated with each row (gene) and column (sample). *Oncoplot* function from maftools, uses ComplexHeatmap Bioconductor package to draw such plots, while providing various customizable options [12]. Oncoplot generated using LAML MAF for top ten mutated genes is shown in Figure 2A. *plotmafSummary* is another function (Figure 2B) which shows overall summary of the cohort in terms of variants per sample and distribution of variants according to variant classification.
- Lollipop plots: Mutations often affects the structure and functions of the protein and it is desirable to represent affected amino acid and conversions on the protein structure. These changes are usually represented by lollipop plots, by plotting protein conversions on to protein structures with underlying protein domains. *lollipopPlot* function takes protein change information and draws a lollipop plot. As an example, lollipop plot for one of the frequently mutated gene in leukemia, DNMT3A, is shown in Figure 2C [13].
- Oncostrip: It is often the case to display a specific set of genes from the whole cohort, to display certain associated characteristics such as mutual exclusiveness. We implemented a function *oncostrip*, which plots given set of genes as a matrix, similar to popular oncoprinter tool on cBioPortal [14]. As an example, two genes NPM1 and RUNX1, which show strong exclusiveness in their mutation pattern, are drawn using function *oncostrip* (Figure 2D).
- Transitions and Transversions: *titv* function classifies each single nucleotide variant into either one of the four types of transversions or two types of transitions, referred by the pyrimdine of the mutated base pair and summarizes them in several ways. This summarized data can be plotted using function *plotTiTv* to display overall distribution of changes (Figure 2E).
- Integrating copynumber and somatic variants: *plotCBSsegments* function integrates copy number data (generated by circular binary segmentation) and somatic variants from MAF files, by mapping them on to copy number segments [15]. As an example, copy number data for TCGA barcode TCGA-AB-3009 along with the mapped somatic variants is shown in Figure 3A. This plot shows two genes NF1 and SUZ12, located on copy number deleted segments.
- Rainfall plots: Cancer genomes, particularly solid tumors are characterized by hyper mutated genomic regions [9, 10]. These regions also referred to as kataegis, can be displayed as ‘rainfall plots’ by plotting inter variant distances on a linearized genomic scale. *rainfallPlot* function draws such plot, and as an example, a rainfall plot for TCGA colon adenocarcioma sample, TCGA-AG-2002-01 is shown in Figure 3B.

**Figure 2.**
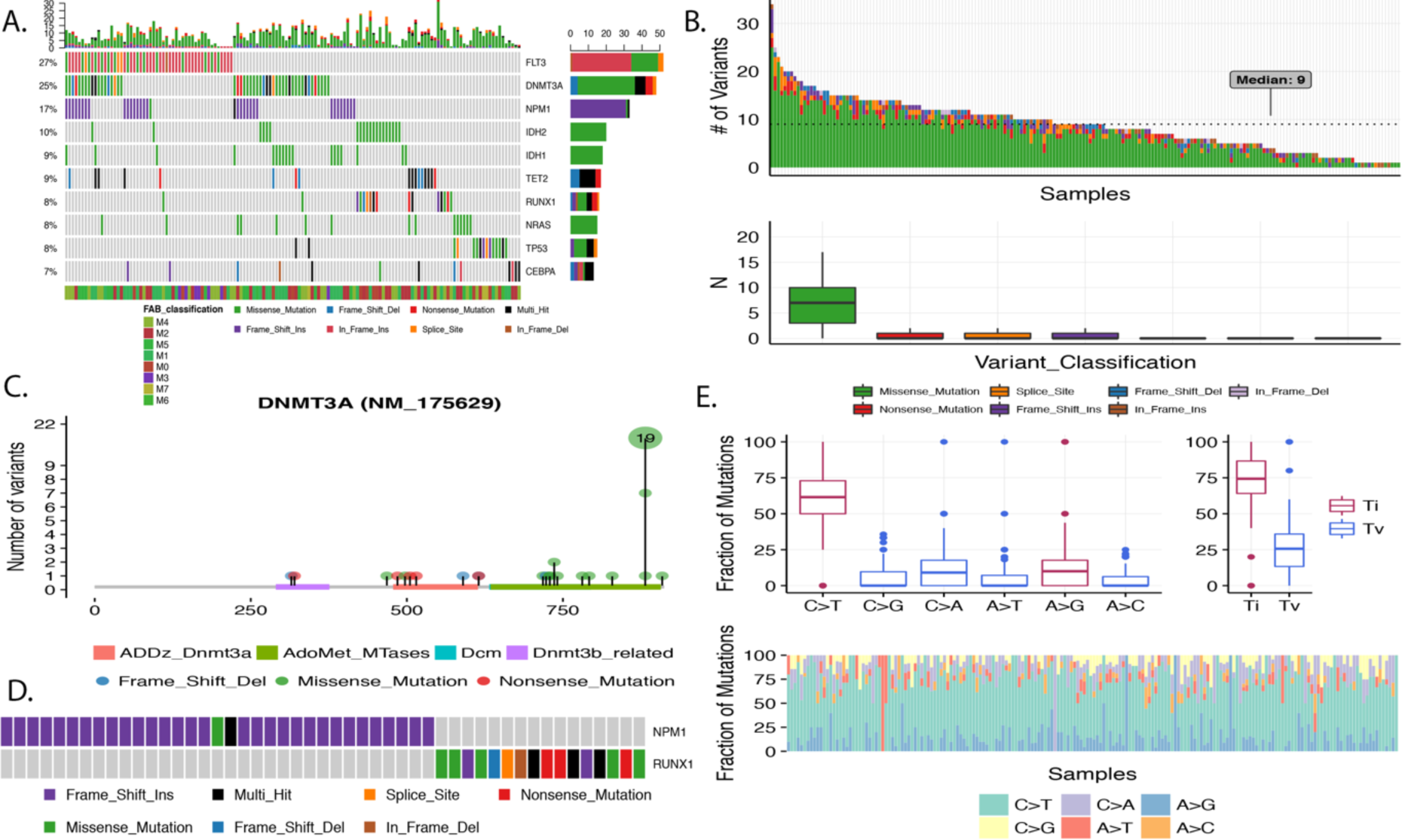
A) Oncoplot depicting top 10 mutated genes sorted and ordered by decreasing frequency. B) MAF summary plot shows a top stacked barplot of variants per sample and a bottom boxplot showing distribution of variants according to variant classification. C) Lollipop plot of DNMT3A. D) Oncostrip displaying two mutually exclusive mutated genes NPM1 and RUNX1. E) Transitions and Transversions–top left boxplot shows overall summary of SNVs classified into six substitution classes; top right boxplot shows distribution of SNVs classified into transitions and transversions. Bottom stacked barplot shows proportion of SNVs per sample classified into six substitution classes.

**Figure 3.**
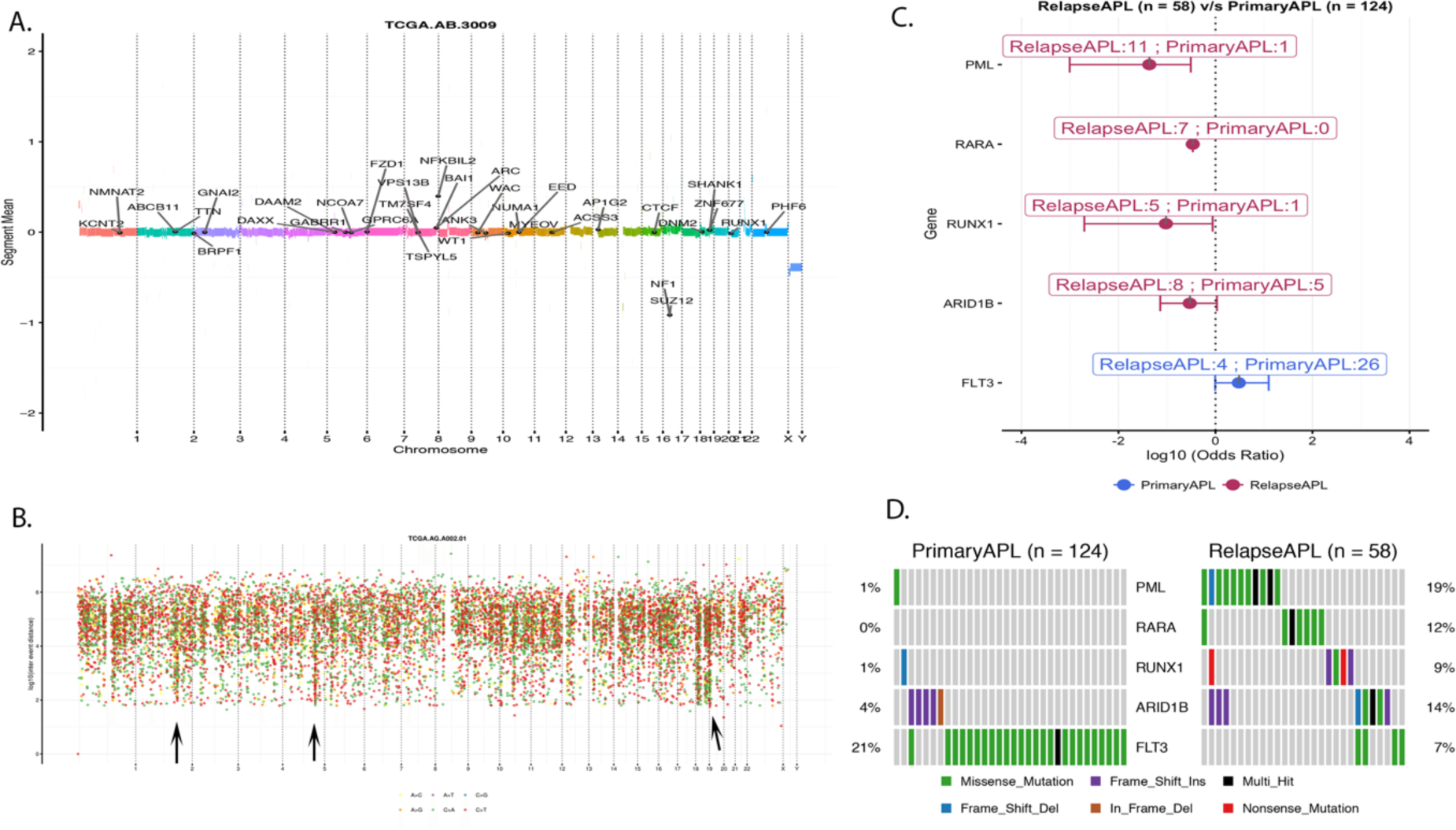
A) Copy number segments for sample TCGA-AB-2009 with the highlighted somatic variants. B) Rainfall plot for colon adenocarcinoma sample TCGA-AG-2002-01. Each dot represents a SNV and are color coded according to six substitution classes. Hyper mutated genomic regions are highlighted with black arrow heads. C) Forest plot for differentially mutated genes between primary APL and Relapse APL. X-axis shows log10 converted odds ratios. Description on each bar shows number of mutated samples in both cohorts. D) Alternative representation of differentially mutated genes as co-oncoplots.

## Analysis

- Differentially mutated genes: Cancers differs from each other by means of mutations in driver genes. This difference is also observable within subtypes of same cancer. Differences between two cohorts can be detected by function *mafCompare*, which performs Fisher’s exact test between two cohorts to detect differentially mutated genes. Recent study by Madan et. al, have shown that patients with relapsed acute promyelocytic leukemia (APL) often harbor therapy induced mutations in PML and RARA gene, which were largely absent during primary disease [16]. Comparing primary and relapsed APL cohorts using *mafCompare*, resulted in five genes (PML, RARA, RUNX1, ARID1B and FLT3) to be differentially mutated (P < 0.05). Results from this comparison are shown as forest plots (*forestPlot*) as well as co-oncoplots (*coOncoplot*). (Figure 3C, 3D).
- Detecting cancer driver genes: Key component of cancer studies is to determine disease associated genes. Often referred to as ‘drivers’, these genes provide selective growth advantage to cancer cells when mutated [2]. To detect such disease associated genes, we built a function *oncodrive*, which is a based on oncodriveCLUST algorithm originally implemented in Python framework [17]. *Oncodrive* takes advantage of the fact that majority of the mutations in oncogenes are clustered around mutational hotspots, whereas mutations on passenger genes are randomly distributed. On TCGA LAML cohort, *oncodrive* was able to detect 11 genes as significantly disease associated genes (Figure 4A).
- Mutually exclusive events: Many disease-causing genes in cancer show strong mutual-exclusiveness in their mutation pattern. *mutExclusive* function performs an exact test on all combinations of genes, to detect a pair showing significant exclusiveness [18]. As an example one of such pair detected in TCGA LAML, NPM1 and RUNX1 are shown as an oncostrip in Figure 2D.
- Pfam domains: Each cancer is characterized by enriched mutations in a particular protein domain [19, 20]. *pfamDomains* function maps protein conversion foci to protein domains and summarizes them according to frequency of mutation. On LAML dataset, most frequently mutated domains include ‘PKC_Like’ (Protein Kinase Catalytic Domain, mutated 54 times across 5 genes), ‘PTZ00435’ (isocitrate dehydrogenase, mutated 38 times across two genes) and ‘AdoMet_MTases’ (S-adenosylmethionine-dependent methyltransferases, mutated 31 times in just one gene-DNMT3A). (Figure 4B). This function is similar to pfam annotation module from Mutational Significance in Cancer (MuSiC) package [8].
- Tumor heterogeneity: It is now well established that tumors are heterogeneous, made up of multiple clones and are constantly evolving [21]. This heterogeneity can be inferred by clustering variants according to their allele frequencies [22, 23]. *inferHeterogeneity* function uses density based finite or infinite (dirichlet process) mixture models, to cluster and classify variants into sub clones [24, 25]. An example of *inferHeterogeneity* is shown on LAML barcode TCGA-AB-2972 (Figure 4C), which clearly shows the presence of two clones, a major clone at a mean variant allele fraction of 0.45 and a minor clone at 0.25 VAF. Options are also included to exclude variants on copy number altered regions.
- Tumor heterogeneity Scores: Mroz et al, have recently shown that the extent of tumor heterogeneity can be measured in terms of numerical values and they introduced MATH (Mutant Allele Tumor heterogeneity) scores; which in short, measures the width of VAF distribution [26]. It has also been shown that tumors with higher MATH scores are prone to poor prognosis and survival [27]. We implemented *math.score* function, which calculates MATH scores based on the distribution of VAFs. Using LAML MAF file, MATH scores for two samples are shown in Figure 4D. Tumor with higher MATH score (sample TCGA-AB-2849; MATH score = 20.59) shows a wider peak, whereas lower MATH score sample (TCAG-AB-2972; MATH score = 11.62) shows a sharp and narrower peak.

**Figure 4.**
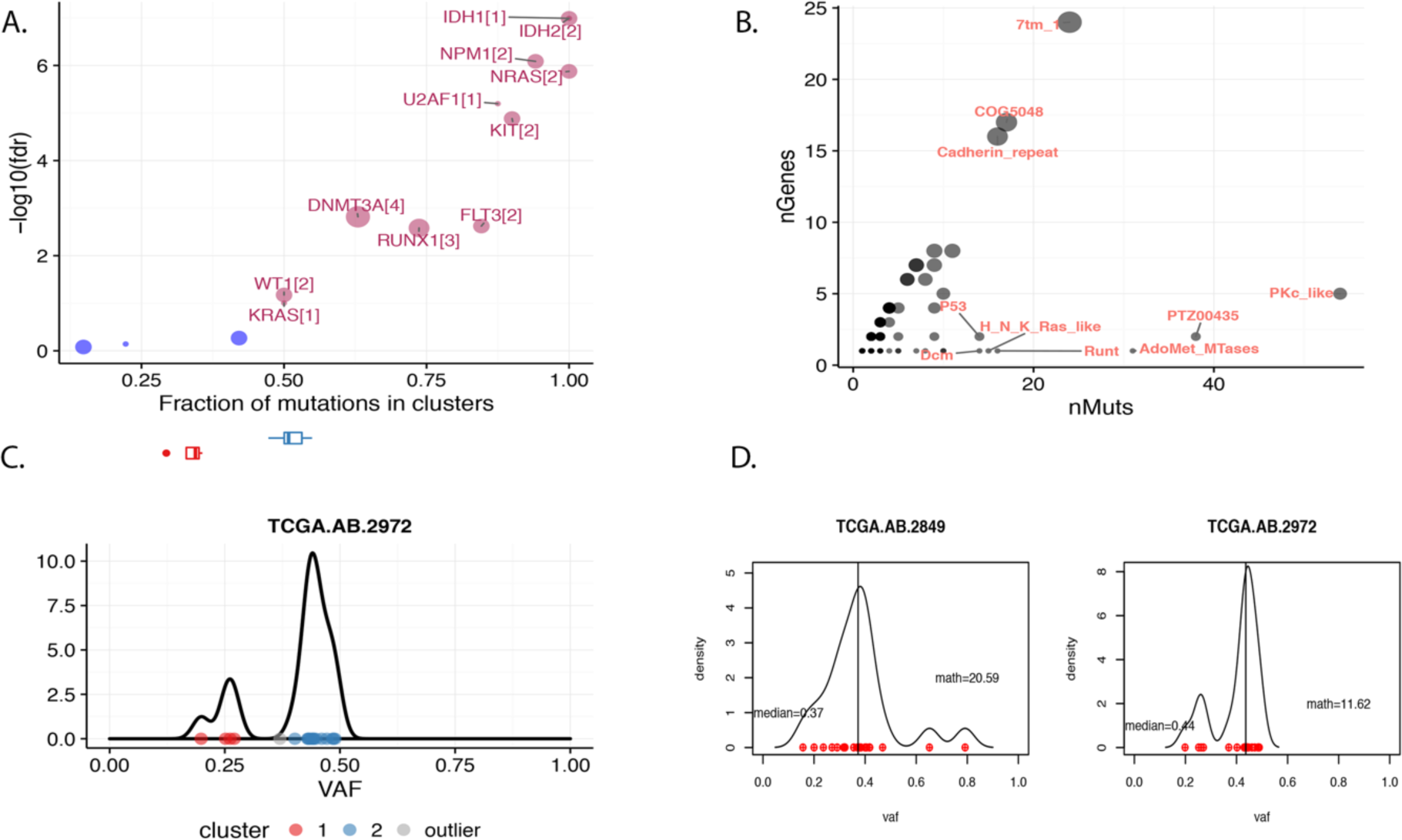
A) Disease associated genes identified in LAML based on positional clustering of variants. Each dot represents a gene and size of the dot represents number clusters (mentioned inside square brackets) within which, a fraction (X-axis) of total variants are accumulated. For example, IDH1 with a cluster score and cluster size of one indicates all of IDH1 mutations are clustered within a single cluster. B) Mutated pfam domains in LAML cohort. X-axis shows number of mutations associated with the domain, Y-axis shows number genes with the mutated domain. Size of each dot is proportional to the number of genes in which the domain is mutated. C) Density plot of VAFs for barcode TCGA-AB-2972. Variants are separated into two clusters. Top horizontal boxplot shows independent distribution of VAFs for each identified cluster. D) Density plot of VAFs for two samples with MATH scores. Each dot represents a variant; vertical bar indicates median value of VAF distribution.

## Variant Annotation and Summarization

Maftools also includes functions for variant annotation and format conversions. Function *oncotate*, takes input variants and annotates them using Broad’s Oncotator web API, and converters them into MAF format [28]. Another function *annovarToMaf* converts annotations generated by popular annotation programs – annovar, into MAF files [29]. Other functions such *subsetMaf*, allows user to filter and subset MAF files on the fly.

## Conclusion

maftools provides a wide range of functions to carry out routinely performed analyses and visualizations in cohort based cancer studies. Maftools was developed while keeping in mind to reduce the burden and hassle of using various software’s/packages, which often requires researcher to change the input data format, and provides a single package on a well-established platform.

